# Metaplasticity: Dark exposure boosts excitability and visual plasticity in adult human cortex

**DOI:** 10.1101/2022.09.04.506561

**Authors:** Seung Hyun Min, Zili Wang, Mengting Chen, Rongjie Hu, Ling Gong, Zhifen He, Xiaoxiao Wang, Robert F. Hess, Jiawei Zhou

**Author notes:** co-first author with SM.

## Abstract

An interlude of dark exposure for about one week is known to shift the excitatory/inhibitory (E/I) balance of the mammalian visual cortex, promoting cortical plasticity across species and accelerating visual recovery in animals that have experienced cortical lesions during development. However, the translational impact of our understanding of dark exposure from animal studies on humans remains elusive. Here, we use magnetic resonance spectroscopy as a probe for E/I balance in the primary visual cortex (V1) to determine the effect of 60-min dark exposure, and measure binocular combination as a behavioral assay to assess visual plasticity in 18 normally sighted human adults (13 females). To induce neuroplastic changes in the observers, 60-min monocular deprivation was performed, which is known to shift sensory eye balance in favor of the previously deprived eye. We report that prior dark exposure strengthens cortical excitability in V1 and boosts visually plasticity in normal adults. Our findings are surprising, given the fact that the interlude is very brief. We present direct evidence that an environmental manipulation that reduces intracortical inhibition can act as a metaplastic facilitator for visual plasticity in adult humans.

## 1 Introduction

Neural plasticity, a defining feature of the nervous system that permits the brain to modify its response and development based on sensory experience, prevails during the critical period after birth and has been believed to irreversibly decline into adulthood. However, molecular and physiological evidence from the animal literature points that visual plasticity in the mammalian adult cortex can be reinstated after the animals experience an interlude of dark exposure [1, 2]. For instance, dark rearing from birth could delay and prolong the critical period across species [3, 4] and reactivate it in adult rats [5] even if animals had experienced a cortical stroke [6], enabling their cortical recovery. Furthermore, amblyopic rats and kittens display a faster recovery in visual functions after a 10-day immersion in the dark [7, 8]. According to animal studies, dark exposure relieves intracortical inhibition and increases cortical plasticity by perturbing the expression of various proteins that are better known as molecular brakes on plasticity, such as chondroitin sulfate proteoglycans [9], perineuronal rats [10] and neurofilament [8], acting as a metaplastic facilitator.

Inspired by the animal literature, human studies have employed short-term monocular deprivation (MD) to induce plasticity in adult humans in the last decade [11]. This protocol mirrors that of animal studies, which assess the susceptibility of the visual cortex to long-term MD as an index of cortical plasticity. The susceptibility in humans to short-term MD is measured as the degree of the shift in ocular dominance that is observed after MD, which favors the deprived eye and is thought to involve homeostatic plasticity that regulates the intracortical excitatory/inhibitory (E/I) balance [12]. Moreover, GABA is decreased in regions within the primary visual cortex (V1) associated with the previously deprived eye [13], suggesting a clear relationship between the perceptual gain of the deprived eye and its reduced GABAergic inhibition in pertinent areas within adult V1.

Using magnetic resonance spectroscopy (MRS), we show that a brief interlude of darkness (analogous to dark rearing in animals) increases the excitability of the visual cortex in humans. Furthermore, we report with supporting behavioral evidence that it boosts visual plasticity in the form of an increased magnitude and longevity of the perceptual gain from the deprived eye after MD. Our findings present the first direct evidence that an environmental manipulation that reduces intracortical inhibition can also act as a metaplastic facilitator for visual plasticity in adult humans, expanding insights into the neurophysiological role of E/I balance in promoting visual plasticity and highlighting clinical applications of metaplasticity.

## 2 Results

### 2.1 Resting concentration of Glx shifts after dark exposure

Eighteen adults with normal vision were tested in this study. Magnetic resonance (MR) spectra were obtained with a MEGA-PRESS sequence [14] from the primary visual cortex (V1) near the calcarine sulcus with a voxel size of 20 × 30 × 20 mm3 (Figure 1A). Each observer completed three MRS sessions. The first session was practice, and its data were excluded. Next, the subjects stayed in the brightly lit lab - where they browsed the web or performed office tasks - for 60 minutes, and then completed the second MRS session with both eyes open. Then, after the subjects completed 60-min dark exposure while both eyes remained open with dark patches on, the third MRS session was performed. Figure 1B illustrates an observer’s spectra before and after 60-min dark exposure. Only the participants who showed similar Glx and GABA concentrations between the first and second sessions were allowed to participant in the study to rule out measurement variability. Glx proxies for cortical excitation and encompasses both glutamine and glutamate, whereas GABA denotes cortical inhibition [15].

**Figure 1:**
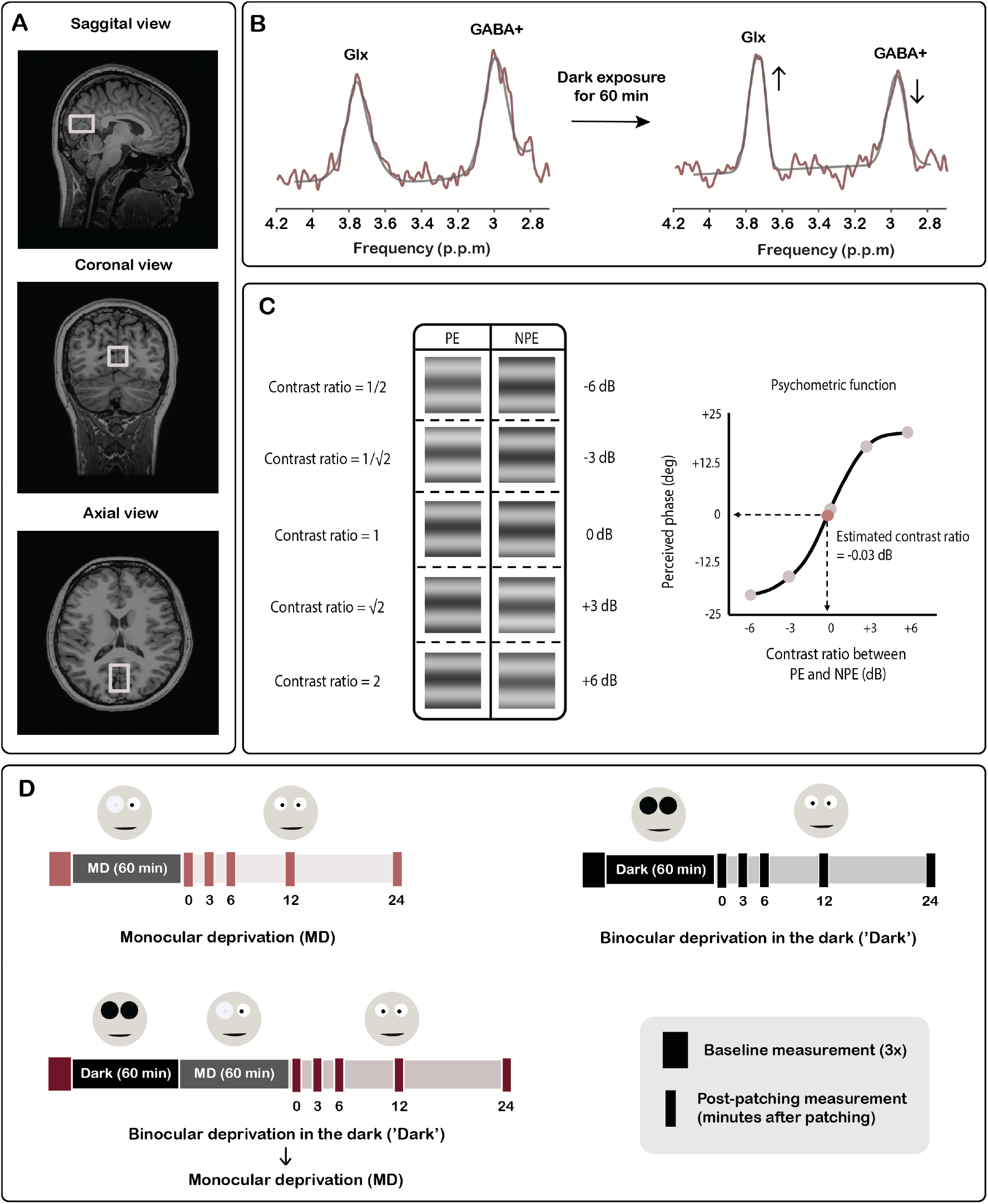
MRS and behavioral experiments. **(A)** Sagittal, coronal and axial views of the representative MRS voxel placement for the primary visual cortex (V1). **(B)** A representative observer’s edited spectra from the visual cortex. Data from two MRS sessions (before and after 60-min dark exposure) were included in data analysis. **(C)** Binocular combination paradigm. Sinusoidal gratings were dichoptically presented at five contrast ratios for baseline measurement (pre-MD) and at three ratios for post measurements (post-MD). Using the data, we fitted a psychometric function to estimate the eye balance. **(D)** Behavioral experiment design. There were three sessions of baseline tests, and then either 60-min MD, 60-min dark exposure followed by 60-min MD (‘Dark -> MD’) or 60-min dark exposure (‘Dark’) alone, and finally post-MD tests at 0, 3, 6, 12 and 24 minutes after patch removal. MD = monocular deprivation of the dominant eye.

Figures 2A-D show the normalized concentrations of Glx and GABA relative to both H_2_0 and creatine before and after 60-min dark exposure. A paired two-sample t-test showed that there is a significant increase in both Glx:H_2_0 and Glx:Creatine between before and after experiencing the darkness (Figure 2A-B; p’s < 0.05 and Cohen’s d > 0.8) but not in the decrease of the GABA concentration (p’s > 0.2, Cohen’s d < 0.5). Lastly, we computed the concentration ratio between Glx and GABA in decibels unit, 20*log(Glx/GABA) referenced to water and Creatine. The ratios were found to significantly increase (p’s < 0.001), suggesting a boost in intracortical excitability in V1. In sum, these results show that a brief dark exposure (60-min) introduces a shift in the concentration of Glx within V1 that does not depend on measurement variability from the MRS scanning procedure.

**Figure 2:**
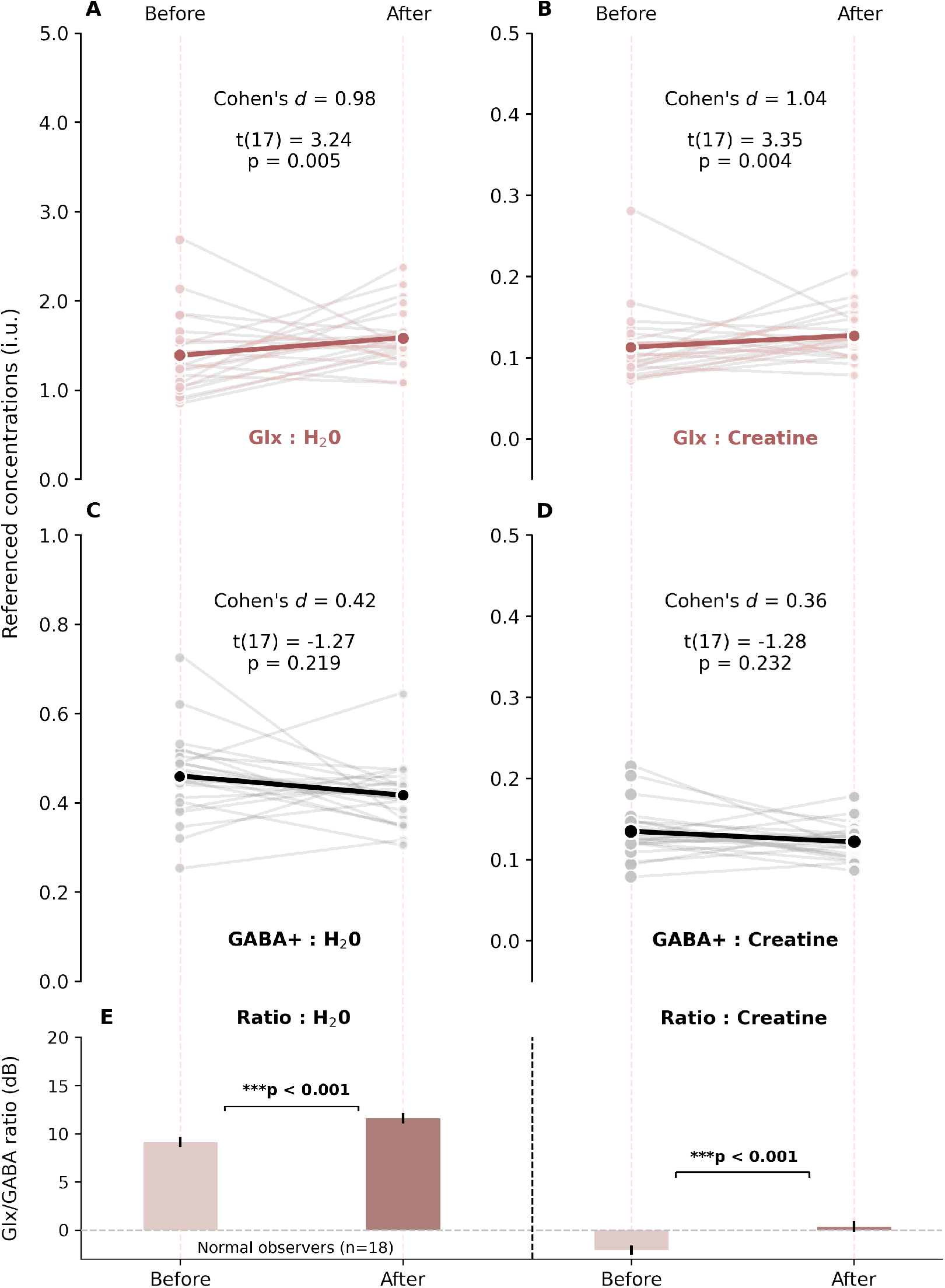
Effects of dark exposure on E/I balance in V1. **(A-D)** Slope charts showing that Glx:H20 and Glx:Creatine significantly increase after dark exposure, where GABA:H20 and GABA:Creatine slightly decrease with no statistical significance. **(E)** A bar graph showing that the E/I ratio as a whole shifted after 60-min dark exposure. Error bars denote standard error and the bar represents the mean of the sample. The grey dashed line represents no shift in E/I ratio, and the peach dashed lines through all panels represent the timepoints of before and after dark exposure.

### 2.2 Perceptual changes of MD are boosted with dark exposure

Binocular balance using a combination paradigm was measured before and after three forms of visual deprivation (Figure 1D; 0 to 24 minutes after patch removal). The first type of deprivation was a 60-min MD. The second type began with a 60-min dark exposure, followed by 60-min MD (Dark -> MD). The last type was an interlude of darkness (‘Dark’) for 60 minutes without subsequent MD. The order of the conditions was randomized for each observer. The data are indicated in Figures 3B-D. In line with previous reports [11, 16], we found that the deprived eye experienced a perceptual increase in combination. This is shown in Figure 3B, which shows changes in eye balance are positive. The deprived eye did not experience a gain after binocular deprivation (‘Dark’) for 60 minutes (see Figure 2D) as the change in eye balance is around 0 (one sample t-test, p’s > 0.05). This suggests that changes due to a period of dark exposure are only effective after MD and are not effective in its own right. Surprisingly, the magnitude of shift in eye balance in Dark -> MD measured at 24-min is comparable to the counterpart in MD at 0-min, indicating that the change is more long-lasting and potent after a brief dark exposure.

**Figure 3:**
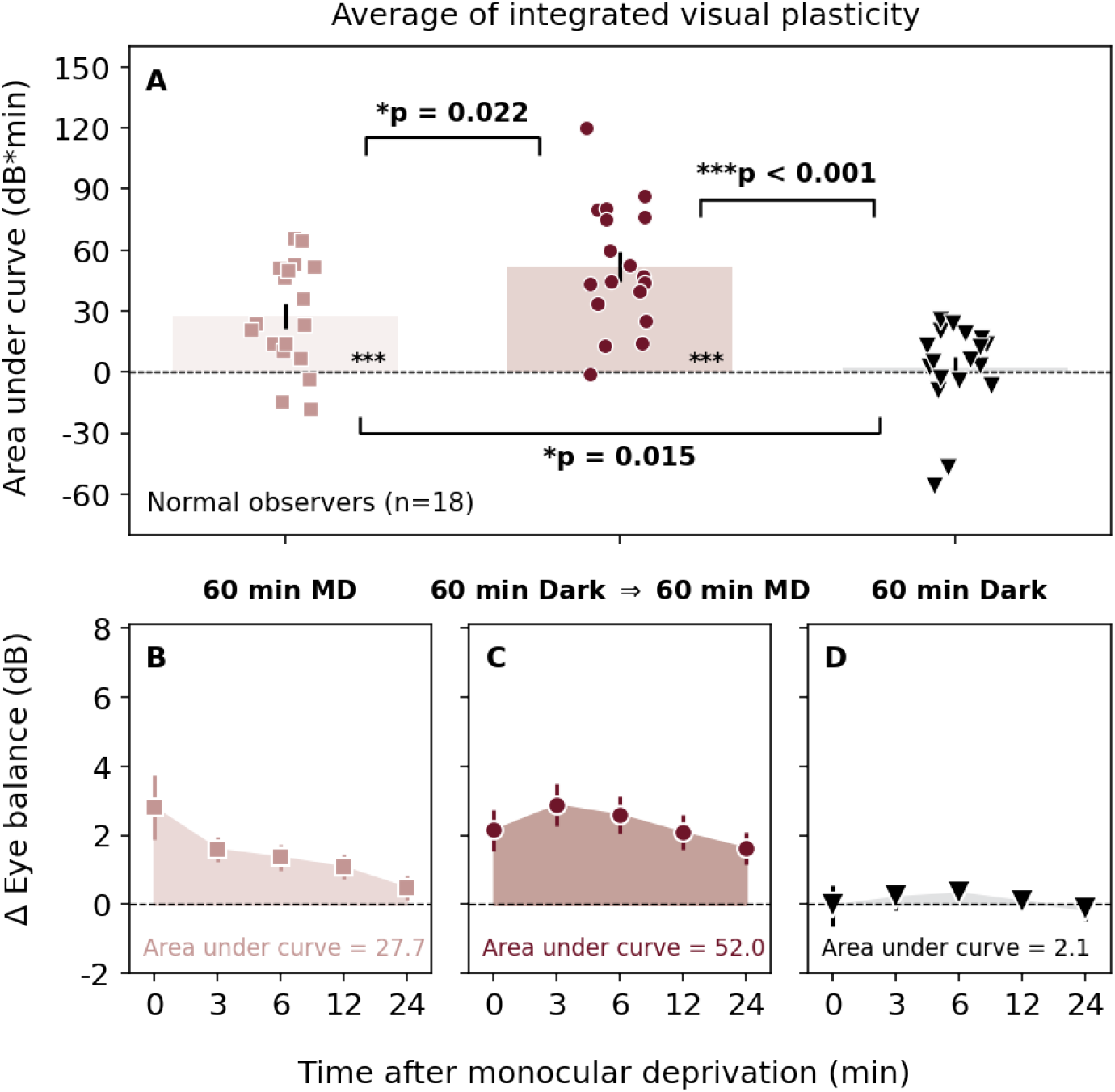
Effects of dark exposure and MD on binocular balance. **(A)** Averaged AUCs across all observers. Asterisks denote a statistical significance based on post-hoc TukeyHSD test or one-sample t-test (*p < 0.05, **p < 0.01, ***p < 0.001). 60-min MD is shown by light red, MD -> Dark by dark red, and 60-min Dark by black. **(B-D)** Changes in binocular balance after short-term monocular deprivation (averaged across all observers). Same colors are used as those in panel A.

In addition, we computed the area under a curve (AUC) of changes in eye balance at various time points after patch removal relative to baseline (Figure 3A). This enabled us to capture both the magnitude and longevity of the induced visual plasticity, and compare it across the three deprivations. Figure 3A indicates that the area is largest in Dark -> MD, followed by MD and the Dark. A one-way repeated measures ANOVA verifies this observation by showing a significant difference in the AUCs across the three conditions (F(2,51) = 16.06, p < 0.001, η^2^ = 0.39). A post-hoc comparison using Tukey HSD (Honestly Significant Difference) test indicated that there was a significant difference in the areal measure between MD and Dark -> MD (p = 0.022), MD and Dark (p = 0.015), and Dark -> MD and Dark (p < 0.001). A one sample t-test showed that the area was significantly different from 0 in MD and Dark -> MD conditions (p’s < 0.001), but not so in Dark.

## 3 Discussion

Together, MRS and behavioral evidence reveals that a brief dark exposure boosts excitability and visual plasticity in adult V1. Importantly, we also provide the first direct evidence that an environmental substrate that boosts cortical excitability in V1 also increases visual plasticity in humans, thereby supporting the widespread belief that the mechanism that underlies reactivating plasticity in humans can be in part due to a shift in E/I balance.

This metaplastic property of darkness has been observed across several species in other forms, such as an extended critical period and boosted visual recovery after artificial induction of amblyopia through long-term MD or cortical damage upon birth [17, 18, 19]. Here, we show for the first time that dark exposure also possesses the metaplastic quality in adult humans by potentiating the neuroplastic changes after short-term MD. Our finding that visual plasticity can be increased after a 60-min interlude of darkness in adult humans is surprising, given the fact that the interlude is relatively brief. In the studies of mice and rats, the interval of up to 10 days has been explored. Our findings indicate that for adult humans there is no need to introduce an interlude of darkness for days or months to induce a metaplastic facilitation, a duration of which could disrupt the circadian rhythm and raise endocrinological consequences beyond the nervous system.

Thus far, our knowledge of increasing cortical plasticity in animals has not yet borne fruits in treating adult humans with cortical disorders, such as amblyopia [20]. Despite clear evidence in animals, a cocktail of approved drugs such as cholinesterase inhibitors [21] that have been shown to facilitate cortical plasticity in mice has yet to demonstrate an improvement of the rate of perceptual learning or visual recovery in humans with amblyopia [22]. Perhaps, the drug dose that has been used in animal studies is much more physiologically potent for animals than what can be provided for humans, failing to elicit its putative effect. Moreover, directly interfering with the neurochemical milieu using drugs raises both ethical and safety issues. Therefore, combined with its disappointing outcome in the recovery of amblyopia, there is limited appeal for implementing pharmacological intervention. The purported mechanism by which these drugs operate to increase cortical plasticity in humans is believed to be through shifting E/I balance in favor of excitation [21, 23]. Our findings in adult humans highlight that a short period of dark exposure can not only tap into the same mechanism but also do so in a less invasive fashion as a metaplastic modulator. Future work will explore whether an interlude of darkness before treatment can be therapeutically used to reinstate plasticity and bring larger perceptual improvements for the population with cortical disorders that impair their vision, such as amblyopia, or other sensory functions.

## 4 Acknowledgment

This study was supported by the National Natural Science Foundation of China (#31970975), the Natural Science Foundation for Distinguished Young Scholars of Zhejiang Province, China (#LR22H120001), and the Project of State Key Laboratory of Ophthalmology, Optometry and Vision Science, Wenzhou Medical University (#J02-20210203) to JZ, the Zhejiang Basic Public Welfare Project (#LGJ20H120001) to ZH, and the Canadian Institutes of Health Research (#125686) and the Natural Sciences and Engineering Research Council of Canada (#228103) grants to RFH. The sponsor or funding organization had no role in the design or conduct of this research. We would like to thank Alex S. Baldwin for giving us the permission to use his program and sharing us the LATEX template (www.github.com/alexsbaldwin/biorxiv-inspired-latex-style).

## 5 Materials and Methods

### 5.1 Participants

Eighteen observers (13 females, mean ± SD of age = 24.1 ± 2.1 years old) with normal or corrected-to-normal vision, including two authors, participated in the study. Using Mile’s test, we established the eye dominance of each subject. The study followed the tenets of the Declaration of Helsinki and was approved by the Ethics Committee of Wenzhou Medical University and the University of Science and Technology of China. All observers were naive to the purpose of the experiment and provided informed consent.

### 5.2 Magnetic Resonance Spectroscopy

#### 5.2.1 Data Acquisition

Magnetic resonance scanning was conducted on a 3.0-Tesla MRI scanner (Discovery MR750, General Electric, USA), with an eight-channel high-resolution radio-frequency head coil within the University of Science and Technology of China. Prior to the scanning procedure, all observers were asked to lie down with eyes open and not fall asleep during scanning. Structural 3D T1-weighted MRI of brain was acquired using T1-3D BRAVO sequence with the following parameters: repetition time (TR)/echo time (TE): 8.16 ms/3.18 ms, flip angle = 12°, 252 axial slices with no gap, matrix: 256 × 256, field of view (FOV): 256 × 256 mm, slice thickness: 1 mm.

The glutamic acid (Glu) and aminobutyric acid (GABA+) were quantitatively measured using the 1H MRS technology. The spectra were acquired with a monosantoid method based on a MEGA-PRESS sequence (Mescher et al., 1998): TE=68ms, TR=2000ms, NEX=8, 256 transients of 4096 data points obtained at 5 kHz. The duration of the scanning was 9 min 20-sec. A 17 ms Gaussian editing pulse was applied at 1.9 (ON) and 7.46 (OFF) ppm. The voxel size was 20 × 30 × 20 mm^3^ at the primary visual cortex (V1) near the calcarine sulcus. The water suppression band of the editing pulses was applied at 4.68 ppm using Chemical Shift Selective saturation (CHESS) imaging [24] and outer volume suppression (OVS) [25]. The very selective suppression (VSS) method in OVS technology was used for the volume of interest (VOI) extra-spatial signal suppression [26].

The ON spectral line was the spectral line that was edited by the spectrum, and the OFF spectral line was the opposite spectral line. During the scanning process of the MEGA-PRESS sequence, an empty sweep period of 8 repetition times (TRs) was required at the beginning of the sequence to allow the signal to reach a steady state during the acquisition. In addition, the reference spectrum containing the water signal was acquired before the ON and OFF spectral line acquisitions. During acquisition of the phase correction reference signal, the CHESS module in the MEGA-PRESS sequence was turned off for 16 TRs.

#### 5.2.2 Data Processing

The quantitative measurement of GABA+ and Glx was completed by editing the MRS spectra using Gannet 3.1.5 (GABA Analysis Toolkit) [26] and inhouse scripts in MATLAB (Mathworks, Natick, MA). All MRS data were analyzed within the .7 format. The water reference signal and the edit spectrum signal were isolated from the MRS data, and the water reference signal was used to correct for the edited spectral signal. The end of the editor spectrum signal was filled with zeros, after which the editor was subjected to a Fourier transform. Then the phase correction was performed on the edit spectrum signal in the frequency domain. The edited spectral signals were aligned to frequencies; the chemical shifts of the Cr signals were aligned to the standard value of 3.02 ppm.

The remaining water signal of the average ON and OFF spectra were isolated from the edited spectral signal, removed, and their baseline corrected. The ON and OFF spectrum were subtracted to produce the DIFF spectrum, from which GABA+ (3 ppm) and Glx (3.8 ppm) signal intensities were modelled using a Gaussian model. The area under the peak of the GABA signal was calculated to obtain the quantitative level of GABA. The areas under the peak of the water signal and Cr signal were used to assess the fitting error of the edit spectrum signal. All the neurochemical signal intensities were considered as the area of the fitted peak(s) and expressed in institutional units (i.u.), using the unsuppressed water and the creatine signal as the internal concentration references.

### 5.3 Behavioral Measurement

#### 5.3.1 Apparatus

The experiments were coded via Matlab and Psychtoolbox, and was conducted on a Macbook Pro laptop. Head-mount goggles (NED Optics Goovis pro, OLED) with a resolution of 1920 × 1080 and a refresh rate of 60 Hz in each eye were used to dichoptically show the visual stimulus. The mean luminance of OLED goggles was 75 cd/m^2^. The equipment was based in Wenzhou Medical University.

#### 5.3.2 Psychophysical Method

In this study, we used a binocular phase combination task to quantify the change in eye balance and induced visual plasticity (Figure 1C). This procedure is described in detail in a previous study [27]. In brief, a sinusoidal grating was shown to each eye with offset phases (−22.5° for one eye, +22.5° for the other eye), a spatial frequency of 0.3 c/deg and a size of 6.6 × 6.6 degrees. When the two eyes contributed equally to binocular version, the darkest strip of the fused grating would be right in the middle of the grating; in other words, the perceived phase would be 0° (the sum of +22.5° and −22.5°). However, if one eye that was shown with +22.5° contributed more to binocular vision, then the fused grating would have a positive, rather than negative, perceived phase. The observers were asked to locate the flanking black reference line next to where they perceived the darkest strip of the fused grating.

The sinusoidal gratings from both eyes were shown at five interocular contrast ratios (dominant eye / non-dominant eye: 1/2, 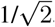, 1/1, 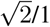, 2/1) for baseline measurement and three interocular contrast ratios (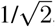, 1, 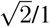) for measurements that were made after the deprivation (i.e., post-patch measurement), and the base contrast was set at 60%. Using the data of perceived phase, a psychometric function was fitted to estimate the contrast ratio that resulted in equal contribution between the eyes [27]. A positive ratio (dB) indicates a stronger dominant (i.e., patched, PE) eye, and the negative ratio a stronger non-dominant (non-patched, NPE) eye. The ratio was computed in decibel units using this formula, 20 * log(contrast of NPE / contrast of PE).

There were eight and five repetitions (i.e., trials) for each interocular contrast ratio during the baseline and post-patch tests, respectively; therefore, there were 80 trials during the baseline measurement (5 interocular contrast ratios × 8 repetitions × 2 configurations) and 30 trials during the post-patching measurement (3 interocular contrast ratios × 5 repetitions × 2 configurations).

#### 5.3.3 Experimental Procedure

To begin with, the subjects completed three sessions of baseline measurement tests (see Figure 1D). Then they underwent a period of visual deprivation (monocular or binocular); there were three types of deprivation, the order of which was randomized for each subject. After the deprivation, they performed post-patching measurement tests at 0, 3, 6, 12 and 24 minutes after the deprivation. The first condition monocularly deprived the dominant eye of the subjects for 60 minutes (MD). In the second deprivation, the subjects underwent an interlude of darkness for 60 minutes with both eyes open and black patches on their eyes. Then, they completed a 60-min MD session (‘Dark -> MD’). During the third condition, the subjects completed 60-min dark exposure without subsequent MD (‘Dark’).

